# Ultra-compliant carbon nanotube stretchable direct bladder interface

**DOI:** 10.1101/580902

**Authors:** Dongxiao Yan, Tim M. Bruns, Yuting Wu, Lauren L. Zimmerman, Chris Stephan, Anne P. Cameron, Euisik Yoon, John P. Seymour

**Affiliations:** Department of Electrical and Engineering and Computer Science, University of Michigan; Department of Biomedical Engineering, University of Michigan; Department of Urology, University of Michigan

**Keywords:** stretchable electronics, biomedical implantable device, bladder control, carbon nanotube, spinal cord injury

## Abstract

The bladder, stomach, intestines, heart, and lungs all move dynamically to achieve their purpose. A long-term implantable device that can attach onto an organ, sense its movement, and deliver current to modify the organ function would be useful in many therapeutic applications. The bladder, for example, is a smooth muscle organ that can suffer from incomplete contractions that result in urinary retention with patients requiring using catheterization. Those affected may benefit from a combination of strain sensor and electrical stimulator to better control bladder emptying. We describe the materials and design of such a device made from thin layer carbon nanotube (CNT) and Ecoflex 00-50 and demonstrate its function with in vivo feline bladders. During bench-top characterization, the resistive and capacitive sensors exhibited reliable output throughout 5,000 stretching cycles under physiology condition. In vivo measurement with piezoresistive device showed a high correlation between sensor resistance and volume. Stimulation driven from Pt-PDMS composite electrodes successfully induced bladder contraction. We present method for reliable connection and packaging of medical grade wire to the CNT device. This work is an important step toward the translation of low-durometer elastomers, stretchable CNT percolation and Pt-PDMS composite, which are ideal for large strain bioelectric applications to sense or modulate dynamic organ states.

## 1. Introduction

Organ disorders can be caused by a great variety of neurologic and non-neurologic events such as peripheral neuropathy, spinal cord injuries, multiple sclerosis, stroke, diabetes, complications of surgery, and simply aging. For one specific organ, bladder dysfunction can have a severe effect on individuals, leading to urinary incontinence, frequency and urgency, and/or incomplete bladder emptying. Among spinal cord injured (SCI) patients who are almost all in urinary retention, regaining bladder emptying function has been reported as one of their highest priorities ^[1][2]^. Patients with SCI also typically cannot sense bladder or bowel fullness and risk having leakage if they get overfilled. In a recent study led by the Craig H. Neilson Foundation, developing an implantable sensor capable of detecting the fullness of the bladder or bowel was stated to be is of critical importance to the SCI patient population^[3]^. Such an implantable sensor might provide input to any existing neuromodulation devices or be part of alternative devices such as direct bladder stimulators. In this work we chose to develop a direct bladder interface integrating strain sensor and stimulation electrode array to address the condition known as detrusor underactivity. The implantable strain sensor alone may also benefit many other conditions or applications.

Patients diagnosed with detrusor underactivity, which causes weak bladder contractions that can lead to retention, who do not respond to conservative treatments like medications and physical therapy are often recommended to use intermittent catheterization to empty their bladders. Although catheterization is a standard treatment, bladder infection rates are high, it is uncomfortable, and can be burdensome. For patients that are unable to self-catheterize for any of these reasons, their only option is the use of indwelling catheters which leads to greater health risks^[4]^. The exact prevalence of detrusor underactivity is unknown, but underactive bladder, a weak bladder muscle in neurologically intact patients, has been estimated as occurring in 20-40% of adult men and women^[5][6]^. Surgical interventions such as implanted devices to stimulate nerves have been a last option for treatment. An implant that applies electrical stimulation to the S3 sacral nerve, known as sacral neuromodulation, has shown the best efficacy for underactive bladder^[7]^, however it does not provide on-demand assistance with bladder emptying and has had poor results in SCI patients and others with neurogenic bladders^[8]^. Electrical stimulation of the ventral roots has been used for effective bladder emptying in SCI patients^[9]^, however its invasive surgery requiring sacral rhizotomy (destruction of all the native nerve roots) has limited its use. In contrast, direct stimulation of the bladder wall holds potential for effective and immediate control over bladder emptying. Prior studies have had challenges with lead migration, recruitment of other pelvic structures or smooth muscle due to current spread, or ineffective emptying^[10]^. Fortunately, advances in stretchable bioelectronics, particularly in materials and fabrication, present new opportunities for direct bladder interface^[11][12][13]^. Furthermore, the use of stretchable bioelectronics also holds promise for detecting the state of the bladder, allowing for closed-loop control over continence and more independence for users.

There are many approaches developed by material scientists and engineers to achieve robust and stretchable electrical connections^[14][15]^, including serpentine metal patterns^[16]^, wavy, buckled^[17]^, origami structures^[18]^, or ultra-thin layer metals^[19]^. Greater strain performance has been demonstrated with conductive polymers^[20]^ and nanowire or nanoparticle percolation networks^[21][22][23]^. Carbon nanotubes (CNTs) are another material for percolation networks, with large aspect ratio structures and many unique physical and chemical properties that have attracted a significant amount of fundamental material science and engineering studies on their synthesis, characterization, and integration. CNTs have inspired scientists to create both flexible microelectronics^[24][25][26]^, and large-strain sensors^[27][28][29]^. For implantable biomedical applications, carbon nanotube percolation networks may hold several key advantages such as inert chemical properties, a large elastic range, and high flexibility^[30]^. Technologies for building layers of CNT include chemical vapor deposition growth^[31]^, spray coating^[32]^, ink-jet printing^[33]^, and roll-to-roll/stamping^[34]^.

Highlighted by the outstanding dynamic range and responsivity, CNT thin layer conductors show great potential as a flexible and stretchable strain detector for wearable electronics applications. To our best knowledge, no examples of applying this material for implantable stretchable electronics have been reported. Several important scientific and engineering issues have yet to be addressed by past research. The reliability and stability of the bare or polymer encapsulated CNT percolation network in physiological conditions is unknown. Failure of the interfacial adhesion and voids in the structure could cause large changes in sensor responsivity or open or shorted circuits. A reliable and biocompatible electrical interconnection also has not yet been demonstrated. Complimentary to CNT percolation structures, one must also use a low-durometer insulator, but others have not yet demonstrated a low durometer silicone attached to a moving organ without tearing either the substrate or the tissue. This work addresses each of these issues. We utilized the low-durometer Ecoflex 00-50 substrate (Smooth-on Inc., shore hardness 00-50), CNT percolation network, a platinum nanoparticle composite, and super-alloy wires to create a functional demonstration of bladder activity sensing and control.

The concept of a direct bladder stimulation array is to access a portion of the bladder surface and suture the perimeter of a stretchable array over the bladder dome and/or posterior wall (Figure 1). Unlike a wrap-around bladder sensor^[13][35]^, this approach requires less surgical time and less risk to the connected nerves and blood vessels, making it potentially more suitable for clinical translation. The built-in sensor detects the bladder wall strain. When the level of strain corresponds to a near-full state, a wireless communicator would indicate this to the patient so a convenient location and time may be found before bladder stimulation and micturition was programmed to begin. Clinical development would also need a wireless IPG/control circuit and a nerve cuff for electrical stimulation that would relax the urethral sphincter during voiding^[36,37]^, which were not developed here. This work moves the field forward by demonstrating improved sensor stability and substrate robustness and discusses the merits of piezocapacitive and piezoresistive sensors in this application.

**Figure 1.**
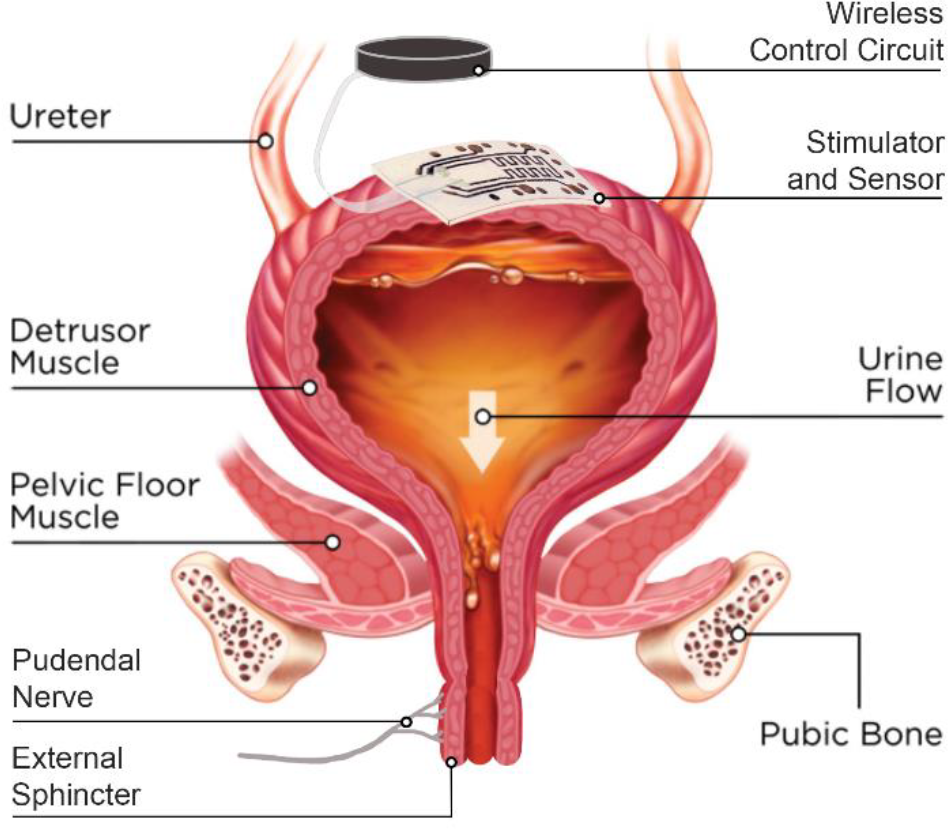
Closed-loop bladder control concept and schematic. This work demonstrated the bladder wall component of this system that includes strain sensing and stimulation. Additional arrays on the dome and/or posterior wall of the bladder would increase stimulation capabilities of the system. Electronic control, wireless telemetry, and a sphincter stimulator would be required for human translation. Identical device and system for men.

## 2. Results and Discussion

We demonstrated two types of organ strain-sensing devices. One deployed a piezoresistive approach with a single CNT layer, and the other deployed capacitive strain sensing with two layers (Figure 2). Each sensor type also has four or more stimulation electrodes for delivering current and causing muscle contractions. The basic principle of the resistive device, as demonstrated in^[38]^, is that the resistance of the CNT resistor increases with increasing strain due to changes in the number of conductive paths in a percolation network. Similarly, the capacitance of two parallel CNT layers increases with increasing strain as the electrode layers expand and the dielectric layer thins during expansion^[39]^. Each sensor had an area of 9.6-mm by 4.8-mm and four stimulation electrodes 2 mm in diameter (Figure 2). The bench-top and *in vivo* performances, mechanical properties, reliability and stability of the device in a physiological environment are discussed in the following sections.

**Figure 2.**
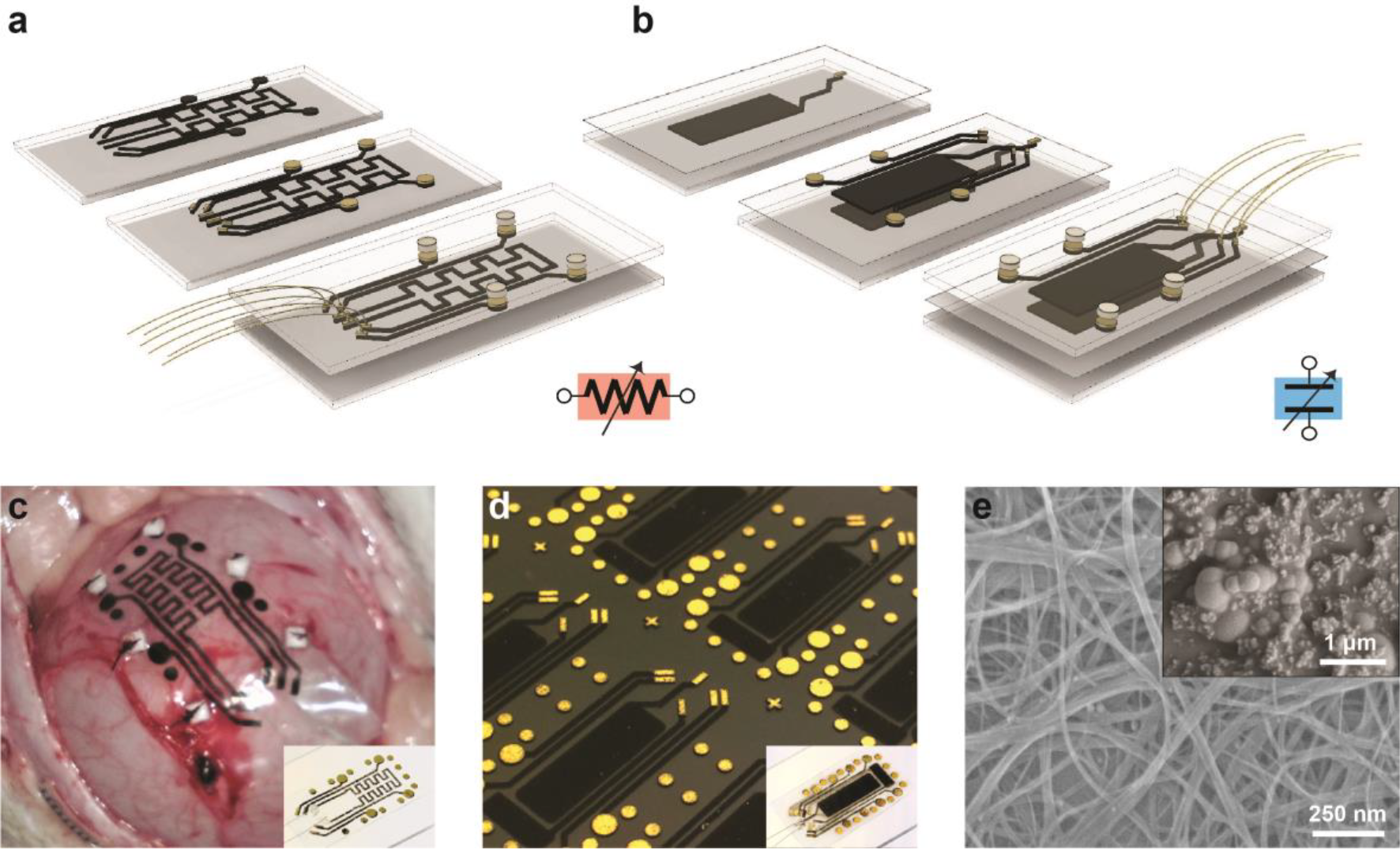
Simplified fabrication process of **a,** resistive-type sensing device with stimulation; **b,** capacitive-type sensing device with stimulation. **c,** A resistive device sutured on feline bladder wall during *in vivo* test. Six 1mm × 1mm white pledgets, a PTFE matrix commonly used for suture reinforcement, are visible on the perimeter. **d,** Snapshot of the capacitive device array on wafer during fabrication. Insets show the single device of each type. **e,** SEM images of the CNT layer and the Pt/PDMS flexible stimulation electrode on top of the CNT/metal layer.

### 2.1. Mechanical properties

Since the bladder expands considerably^[40,41]^, any implant attached on the bladder wall must induce only a negligible amount of force and minimally alter the system mechanics. To determine the bulk material of the device, we conducted elasticity measurements of feline and porcine bladder samples and compared this to PDMS (Sylgard 184) prepared at a 20:1 ratio and Ecoflex 00-50 (Figure 7a). All the samples were 2cm long with a cross section of 3mm by 1.2cm. The total tensile force applied horizontally on one end of the tissue as a function of tissue expansion was recorded. PDMS (Sylgard 184), as one of the most commonly used soft elastic silicone materials, was far too stiff at 100µm thickness, while Ecoflex 00-50 has a significantly lager elastic range and 1/16 the stiffness of PDMS prepared at a 20:1 ratio. The thickness of the Ecoflex substrate was empirically tested from 100µm to 2mm in thickness. We selected a thickness of 300 µm as it did not tear at 120% strain and offered little resistance relative to thicker substrates. Despite its industrial-grade manufacturing, several studies have shown promising results for the biocompatibility of Ecoflex silicone ^[42][43]^. When a 300-µm Ecoflex substrate was attached to a feline bladder wall and the bladder was fully expanded, we observed no deformation in the bladder’s curvature (Figure 7a).

Initially when we attached either PDMS or Ecoflex substrates onto an *ex vivo* porcine bladder with medical suture, local tearing in the substrate was evident after 2-5 cycles because of the high local stress around each suture attachment location. Medical grade cyanoacrylate or other glues may help secure a substrate to the bladder with less local deformation, but it would also impede current flow at the electrode-tissue interface. Instead, we applied commercially available pledgets (small PTFE pads; 1.5 × 1.5 mm) at predefined locations around the perimeter. Each location was marked with a small CNT/Au pattern (Figure 2d) and six pledgets were glued using a drop of Ecoflex 00-50. This technique eliminated tears in the substrate when attached with suture to feline or porcine bladders. One *ex vivo* porcine bladder experiment and four *in vivo* feline experiments with around ten full-strain cycles each were conducted using devices with attached pledgets, and no visible tearing around suture sites occurred.

### 2.2. Device functionality and repeatability during strain cycle

We conducted bench-top and *in vivo* experiments to evaluate the performance of the bladder strain sensor. Both piezoresistive and piezocapacitive devices were immersed in PBS solution heated to body temperature, clamped at each end, and repeatedly stretched from 0% to 100% strain (setup described in Supp. Figure 3). The impedance of each sensor type was recorded for 5,000 cycles at this condition. Before any high-strain use, both device types needed to go through five to twenty pre-stretching cycles to stabilize and reorient the percolation network. The resistance changes during this process is shown in Figure 3 for an example device. A similar need for pre-stretching cycles was reported in previous studies^[29][44]^.

**Figure 3.**
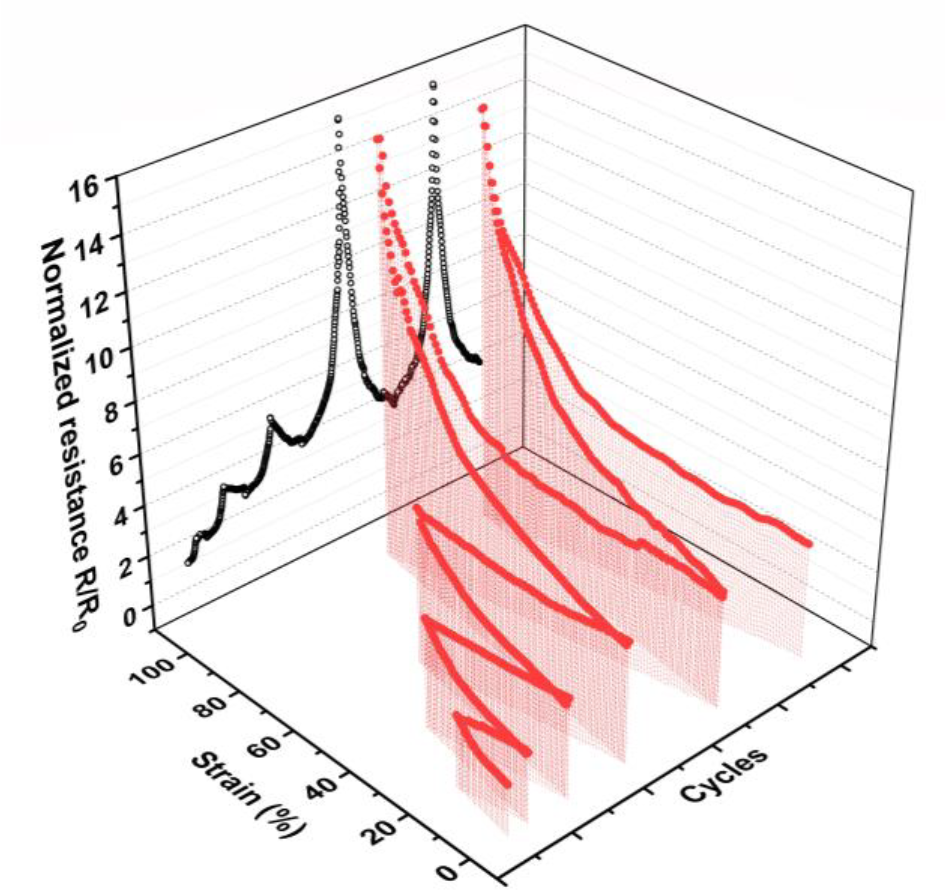
Pre-stretching a CNT conductor. Normalized resistance of a 500-µm wide conductive trace made of carbon nanotube layer (2.9μm thick) during “pre-stretching” cycles. Stable resistance was achieved between the 4^th^ and 5^th^ cycles here.

#### Capacitive and resistive CNT sensors for strain detection

All devices were strained from 0 to 100% in the longitudinal direction. Equivalence to a 100% longitudinal strain with no transverse constraint was estimated to be approximately 78% biaxial strain (ΔS/S, where S is the area) based on total area change. The capacitive devices measured an increase in strain from capacitance change due to a combination of two effects, decrease of the dielectric layer thickness and increase of the electrodes effective surface area from stretching. Four capacitive devices were tested during the bench-top experiments and demonstrated good repeatability (Figure 4b). Although a variance of absolute capacitance values at the free state was observed, the normalized capacitance showed very similar and repeatable strain response among devices. Sensor responsivity was calculated as the input-output gain, ΔZ/Δε (Figure 4a). Despite variances introduced by fabrication, the device can be easily calibrated by dividing the zero-strain impedance. The relative capacitance, C/C_0_, increased from 1.0 to 2.13 ± 0.20 (N=4) in response to the same range of longitudinal strain.

**Figure 4.**
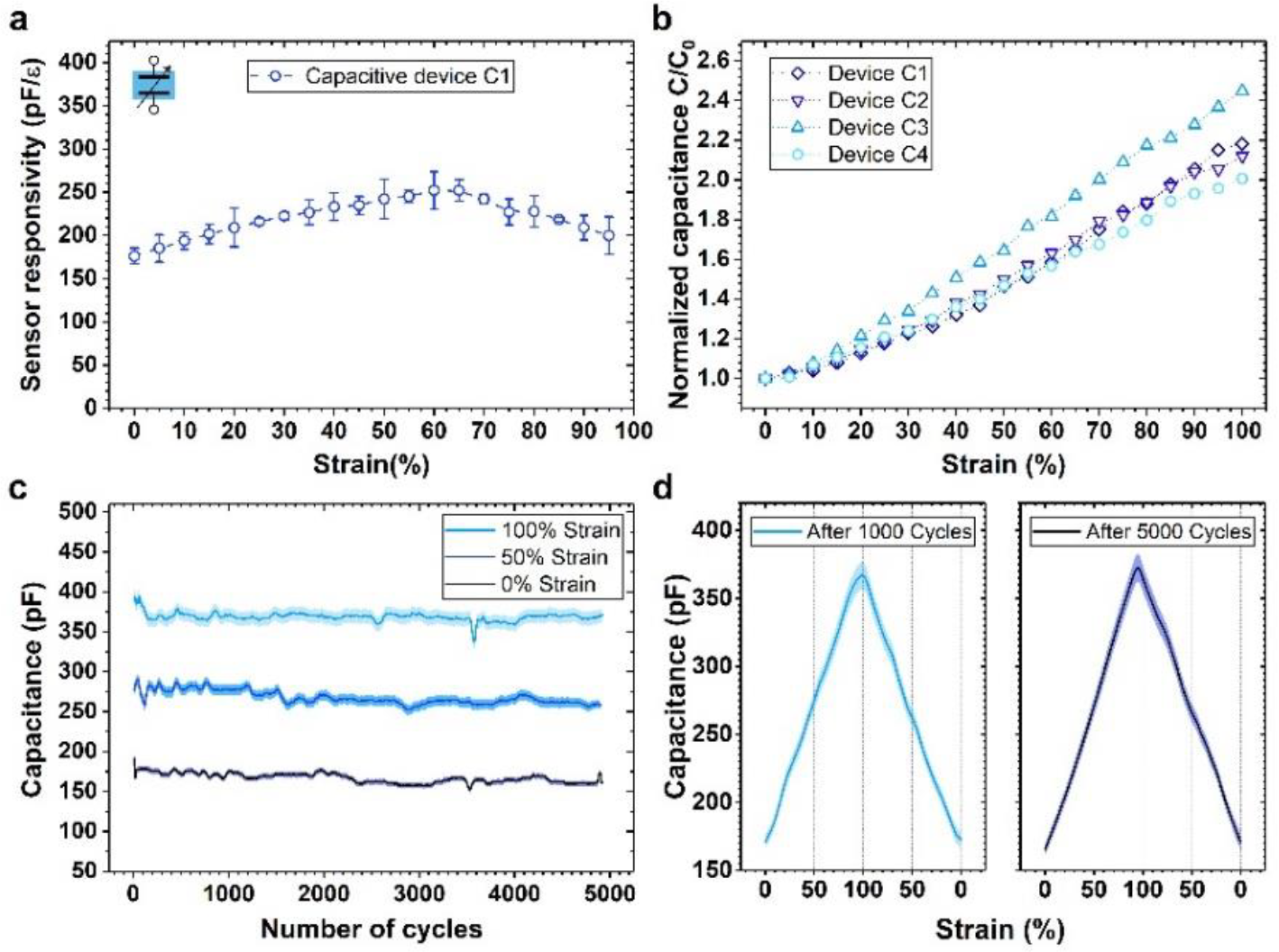
Performance of piezocapacitive sensor. **a,** Capacitive device responsivity to uniaxial strain, ε (longitudinal). **b,** Normalized sensitivity or gauge factor of 4 capacitive devices. **c,** Sensor stability throughout 5,000 stretching cycles at three strain points, 0%, 50% and 100% of each cycle. Shaded area indicates the standard deviation (S.D.) for ± 5 cycles. **d,** Continuous recordings of the sensor response after the 1,000^th^ and 5,000^th^ cycles. Shaded area indicates the S.D. for ± 5 cycles.

**Figure 5.**
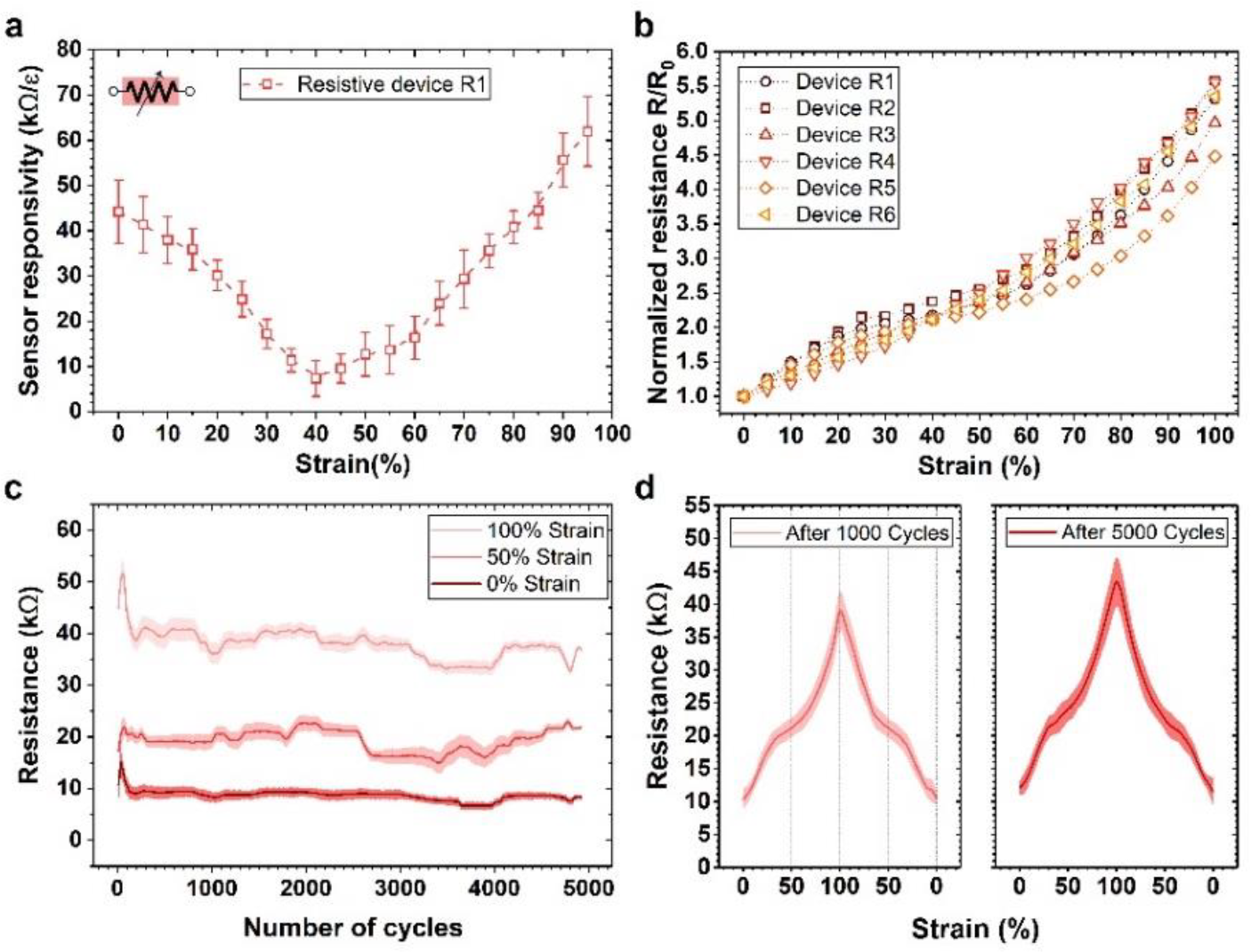
Performance of a piezoresistive sensor. **a,** Resistive device responsivity to uniaxial strain, ε (longitudinal). **b,** Normalized sensitivity or gauge factor of 6 resistive devices. **c,** Sensor stability throughout 5,000 stretching cycles at three strain points, 0%, 50% and 100% of each cycle. Shaded area indicates the S.D. for ± 5 cycles. **d,** Continuous recordings of the sensor response at the 1,000^th^ and 5,000^th^ cycles. Shaded area indicates the S.D. for ± 5 cycles.

As for the resistive devices, during the expansion and contraction of the substrate, the tensile and compressive stress induced a re-orientation process in the CNT network, where each individual CNT repositioned to adapt the macroscopic strain. As a consequence of this dynamic configuration, both the conductance and the quantities of the conductive pathway within the CNT resistor changed with strain, which resulted in a non-linear change in the resistance. Comparing with conventional MEMS piezoresistive materials such as heavily doped silicon^[45]^, polysilicon^[46]^, and so on, the obtained CNT network exhibited a much larger dynamic range while maintaining a large gauge factor. Similar to the capacitive devices, the normalized resistance R/R_0_ can be used as a reliable measurement of strain regardless of absolute value variance among devices.

Bench-top testing showed a reversible and repeatable piezoresistive effect. The relative resistance, R/R_0_, of the CNT sensors increased from 1.0 to 5.1±0.18 (N=6) in response to longitudinal strain from 0% to 100%. The piezoresistive type sensor exhibited higher sensitivity albeit not as linear as the capacitive response. Neither measurements showed a significant shift in the impedance vs. strain curve after 5,000 cycles (Figure 4d and 5d), suggesting that further cycling could be achieved. The resistive devices also exhibited more fluctuation than the capacitive devices in the absolute impedance value during cycling. The reason for the drift in impedance was likely a long-term drift of the CNT network configuration at micro-scale, while the capacitance, or total effective electrode surface area and dielectric layer thickness, was less sensitive to the network configuration. Both types of the devices showed a high overall responsivity at 350.5Ω and 2.1pF per percentage strain, but the former is significantly easier to measure.

#### Stability in physiological environment

It is a highly critical factor for any implantable device to remain stable in a physiological environment. To examine this concern, we collected a series of sensor output data while three devices were immersed in saline solution at body temperature at different time points including immediate, 10-hour, 24-hour, 48-hour and 1-week post immersion. Next, the devices were kept soaked while being stretched under 0% to 100% longitudinal strain for 1,000 cycles. Then the devices remained in the same condition for another 3-weeks (Figure 6). Except for the resistance measurement after 1,000 strain cycles post-1-week immersion, fluctuations of resistance were no larger than ±27% and of capacitance were no larger than ±23% observed during soaking. This is an important first step toward demonstrating the stability of both sensor types made from encapsulated CNT networks and the wire to CNT connection. Although stretchable CNT networks have been utilized as a conductor or semiconductor material for many wearable electronics applications, this is the first successful demonstration of a CNT nanosheet-based strain sensing device in a solution environment for potential implantable medical applications. Note that the electrical stimulation interconnects were also made of the same width and thickness (0.5 mm wide, 2.90 µm thick) and thus are expected to have the same stability.

**Figure 6.**
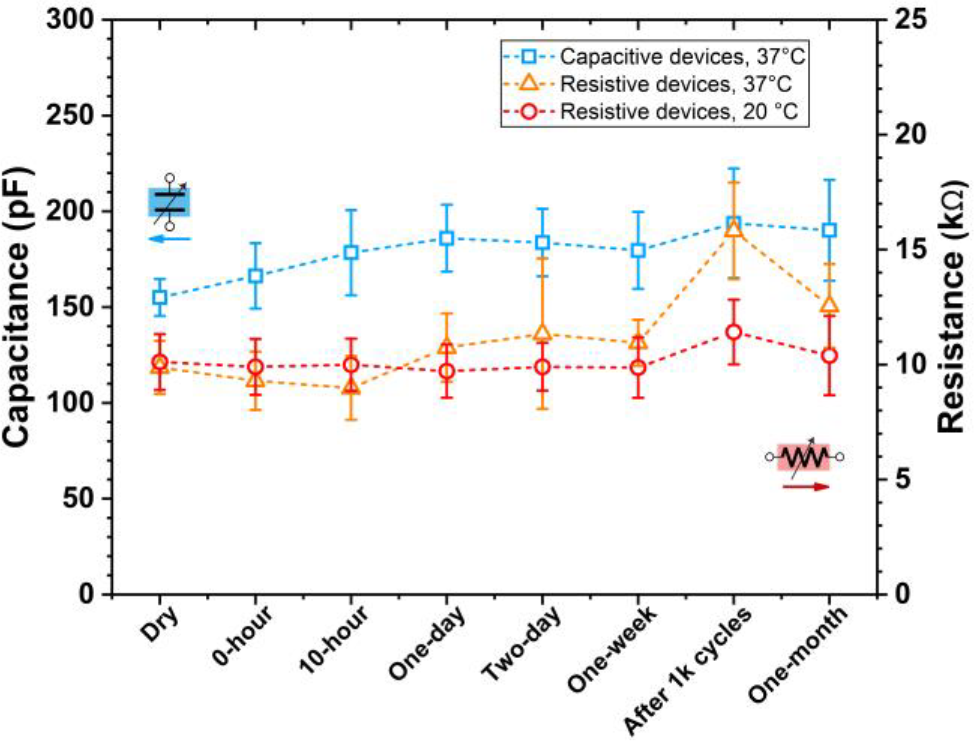
Sensor stability test under physiological conditions. Both resistive and capacitive device were immersed in PBS solution at 37℃. The plot shows the resistance/capacitance change at different time points. N=3 for each.

**Figure 7.**
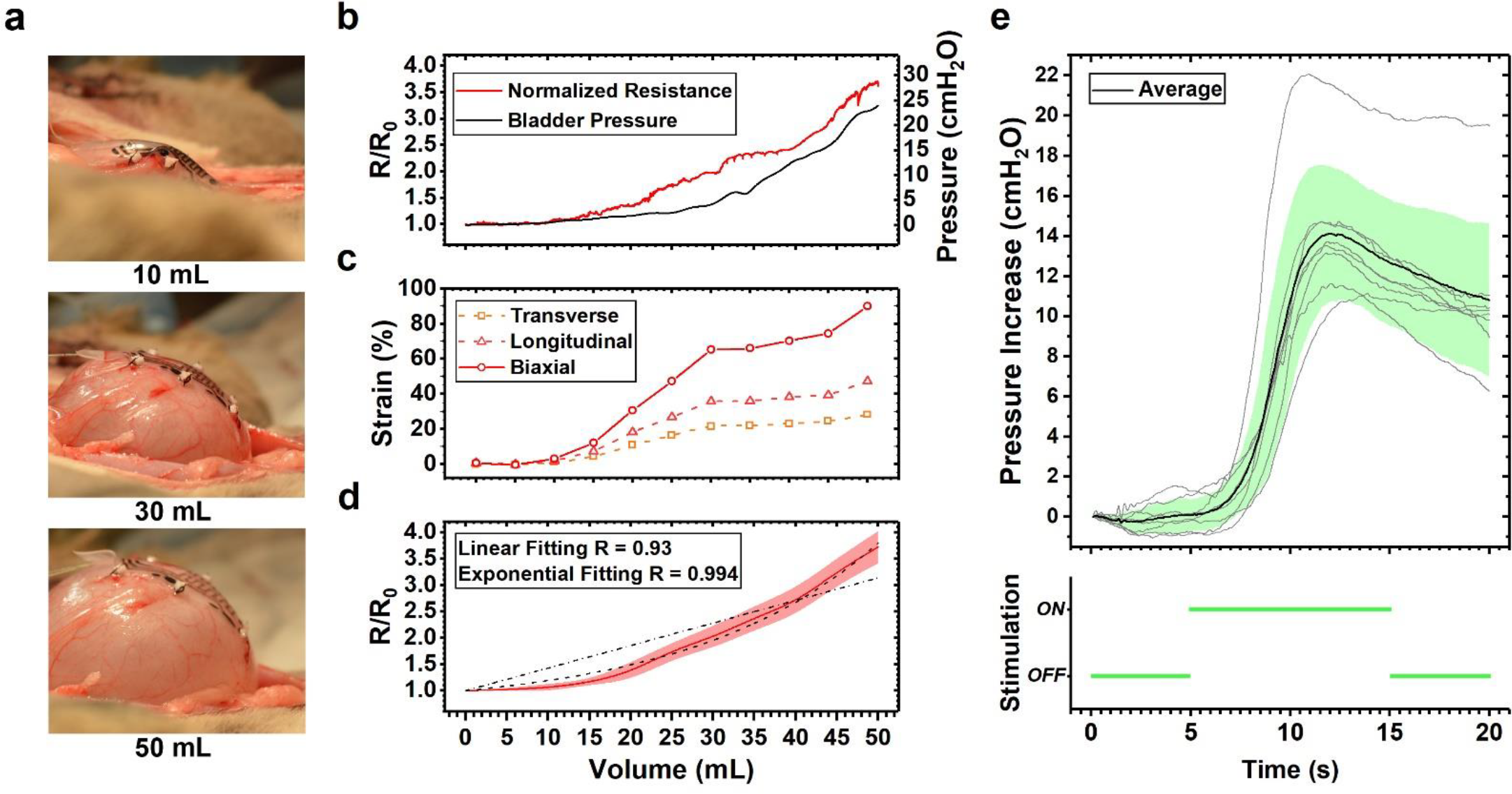
*In vivo* sensor performance and stimulation induced contraction. **a,** Side views of device sutured on a feline bladder with 0, 30 and 50mL liquid filled. Note the attached device does not create noticeable deformation of the bladder. **b,** CNT piezoresistor sensor output responding to the bladder volume during a fill to 50 mL with the accompanying bladder pressure. **c,** Longitudinal, transverse and biaxial strain values of the device during the same trial using image analysis of top-down view. **d,** Mean sensor output across nine bladder filling trials in one animal 3 (S.D. shaded in red). Linear and exponential fit relative to the delivered volume are shown with dashed lines. **e,** Stimulation-induced bladder contractions. All 8 stimulation trials (light gray) overlaid with mean (black) and S.D. (light green) response.

#### Capacitive vs. Resistive Sensing Modes for Bladder Stimulation

Based on our *in vivo* experiments discussed below, the biaxial range may be anywhere from around 45% to 90% strain in practice. This is approximately equivalent to a uniaxial strain of 54% to 110%. For the piezorestive device, the worst drift observed was between the 0-hour measurement and after 1,000 cycles, but after soaking 1-week. The drift at this point was 59%, but then recovered to 27% higher than the initial value after 1-month total soaking. For the piezocapacitor, the drift over the same period was 15%. Percent variance after 1,000 stretching cycles and 1-month soaking for the resistive and capacitive sensors was 27% and 23%, respectively (N=3 devices). However, a very critical difference between the two is the measurement circuit. A low-noise detection circuit for the resistor could be made with a simple voltage-divider but accurately measuring a difference of 2.1 pF per 1% strain difference is a more complex and sensitive circuit. In the latter case, even parasitic capacitance in the lead body and connector to the electronics housing could add significant drift to the total capacitance.

The required accuracy and drift limits for a useful implantable sensor is not yet clearly defined. The simplest requirement is the detection of the point when the bladder is nearing its full state, which ideally is well before full to allow a patient time and convenience to manage the emptying of their bladder. In prior work the correlation between an implantable pressure sensor and a catheter pressure sensor was on average better than r=0.90. As we show below, we too have excellent correlation with resistance and volume, but that does not account for long-term drift. This data taken as a whole suggests that a resistive device is likely to be an easier and more robust sensor and circuit to implement. Future work could further improve on our resistor performance by defining an optimal pre-cycling and pre-soaking regimen before chronic implantation. For these reasons, the *in vivo* measurements were taken using the resistive type devices.

### 2.3. *In vivo* validation including biaxial strain

We tested resistive devices during four anesthetized feline experiments. The devices were first sutured onto the bladder while the bladder was at approximately 1/3 of the bladder capacity. 8 to 14 fill-empty cycles were conducted in each experiment, with the devices stretched up to the limit of the bladder volume at which leakage occurred. Table S1 shows the estimated biaxial strain among all four animals at their maximum bladder capacity. Maximum strain was limited by bladder leakage around the inserted catheter, and somewhat by placement, so variability in the orthogonal and biaxial strain was considerable. One representative trial illustrates the correlation of resistance, strain, and pressure to volume which was controlled with an infusion pump (Figure 7b). The strain, measured from the video footage of this same experiment (Animal 3), corresponded to approximately 49.6% longitudinal and 30.2% transverse strain with no mechanical or electrical failure (Figure 7c). Furthermore, we show the repeatability of normalized resistance versus volume for this same animal (N=9 fill cycles, Figure 7d). Resistance is a measure of strain in that portion of the bladder wall, especially biaxial strain (∆S/S, where S is the area) of the device, and not necessarily linear with volume. The linear correlation linear coefficient for the mean of all cycles was measured to be 0.93 (9 trials in one feline), and could be greater than 0.99 when fit to an exponential equation (Figure 7d). Pressure is not linear to the bladder volume. As a point of comparison, implantable pressure sensors implanted into the wall of the bladder had an average correlation coefficient of about 0.90^[47,48]^.

In one terminal experiment, we pushed the bladder capacity beyond the point of visible leakage at the urethra in an effort to understand the limits of the device and sutures at the last trial. Electrical failure was observed in this trial before any tearing was observed at the bladder by the tissue or the device. This failure point occurred when the observed biaxial strain was above 133% (corresponding to 64% and 53%, longitudinal and transverse, respectively).

### 2.4. Stimulation electrodes and in vivo bladder excitation

CNT can provide efficient charge transfer to tissue mainly because of its exceptionally high surface area^[49]^. However, given the potential cytotoxicity of CNT when directly released into tissue, a coating over the CNT has advantages^[50]^. Furthermore, we desired to limit the strain on the electrode by integrating a stiffer elastomer with a mixture of Pt nanoparticles and Sylgard 184. To improve the charge capacity of the device we utilized a nanocomposite previously demonstrated on an all-PDMS device and having gold interconnection^[51]^. Our Pt-PDMS nanocomposite was screen patterned (2-mm diameter and approximately 150-µm tall) into each electrode recess patterned on the encapsulating Ecoflex layer connecting to the CNT and Ti/Pt/Au layer on the bottom. Whether Pt-PDMS or CNT alone is a superior material is outside the scope of this report, but future studies should consider the biostability and toxicity of CNT as a direct stimulation material.

A four-point test structure was made to measure the contact resistance between the conductive composite and the mental/CNT electrodes. The CNT resistance (2.90±0.28μm thick controlled by the total mass of CNT deposited and 500-µm wide) was measured to be 0.9-5.5 kΩ as-deposited and 4.8-19.4 kΩ after pre-straining cycles. The contact resistance was lower than 50 Ω. The platinum-silicone composite turned out to have moderately low impedance (Figure S4, Z_s_=77±64 Ω∙cm^2^ at 1kHz). The variation was caused by the non-uniform distribution of platinum microparticles inside the PDMS mesh. Unlike other demonstrations of the Pt-PDMS nanocomposite, we used reactive ion plasma etching to remove a thin layer of the top PDMS so that the surface of the embedded platinum particles could be exposed, which reduced the resistivity by five times. Electrode impedance was scanned from 10 Hz to 10kHz in saline solution in a three-electrode system. Cyclic voltammetry was recorded in 0.15M sulfuric acid over the range of −0.3V to +1.1V versus Ag/AgCl electrode (scan rate 50mV/s). The composite showed 4.39±2.64 mC/cm^2^ (Figure S4) cathodic charge storage capability, which was almost order of magnitude higher than a smooth platinum electrode^[52]^.

Electrical energy consumed on all the 4 electrodes during stimulation event can be calculated using the parameters of the bipolar pulses and the average impedance of the electrodes. For human application, a typical stimulation current and duration may be 10mA, 300μs pulse width and 40Hz for 3 seconds^[10]^. To further reduce the stimulation driving voltage to around 10V, one can increase our CNT trace thickness to 10 μm and width to 1mm. Adding the power consumption of a typical cardioverter defibrillator monitoring system as an estimation^[53]^, the device with a 350mAh commercially available primary lithium battery for medical implant (EaglePicher LTC-3PN-M1), assuming 8 bladder empty per day, could potentially operate for over 7 years.

Charge injection capacity and impedance as a function of the device strain did not change much because the PDMS mixed with high weight proportion of Pt micro-particles was much stiffer then the surrounding Ecoflex 00-50. According to a COMSOL simulation (Figure S5), for our particular geometries the expected maximum strain of the Pt-PDMS composite was below 5%. This may be an important advantage since stimulation for this application will only occur near the point of maximum strain and thus a Pt-Ecoflex composite would have been significantly higher electrical resistance and limited current flow.

During an *in vivo* test, bipolar electrical stimulation was applied through the stretchable electrode on a piezoresistive device, using stimulation parameters of 33-50Hz, 3ms pulse width and 1.25-2.5mA. Before testing on the bladder, simply placing the device directly on the abdominal muscle allowed for robust recruitment of the somatic muscle during stimulation (Supplemental Video 1). Next, the device was placed on the exposed bladder before being sutured in place. After the bladder was filled to a near-full volume, ten-second epochs of stimulation successfully induced bladder contractions, with increases of 10.5-21.9 cm H_2_O observed (Figure 7e) from a baseline of about 15 cm H_2_O. The pressure peaked at 6.8±0.6s after initiation of stimulation. Effective bladder emptying in similar anesthetized feline experimental preparations has been reported at 37 cm H_2_O ^[54]^, suggesting that the bladder responses driven here are capable of driving micturition. Larger contractions, of 50-80 cm H_2_O ^[55]^ and above 100 cm H_2_O ^[56]^ have been reported for non-anesthetized animals. Future experiments will explore the optimal geometry to create robust bladder contractions. As the human bladder holds ten times the volume of feline bladders, there is sufficient area for a larger device or multiple devices. Furthermore, we chose a small rectangle for the simplicity of suturing in these studies, but the fabrication technology allows any arbitrary geometry such as a star shapeto provide more access to multiple and specific locations on the bladder.

### 2.5. Packaging

The implantable application of CNT-based stretchable electronics was previously limited by two fundamental engineering concerns—the reliability of insulation and the robustness of wire-bonding. In early proof-of-concept studies^[39,57]^, the wire connection from the CNT layer to an external circuit was achieved by conductive tape and a clamp, or wires were not allowed to stretch. However, in practical use by a surgeon, any device must perform well with noticeable strain on the leads and obviously have well insulated solder joints. To address this issue, we developed a novel, robust, and low impedance electrical interface from a medical grade super-alloy, MP35N (Fort Wayne Metals) to the CNT layer. A 20 nm thin layer of Ti was first evaporated on the CNT layer to form a low resistance metal seeding at the wire-bonding region^[58]^. Secondly, a 50nm Pt layer was evaporated above the Ti layer as a diffusion barrier followed by a 300nm thick Au layer on top to provide a strong and low resistance bonding interface. Next the MP35N alloy wire was bonded to the Au surface by applying silver paste (TED PELLA. Inc. 16040-30) and low temperature annealing at 120°C. Finally, the wire bonding spot was further encapsulated and secured by a thin layer of Ecoflex spun on top.

The built-in strain relief of this technique was surprisingly robust. The wires themselves could be tugged (Supp. Fig. 2) and there was no measured change in resistance even though the local stress to the bond pads was very high. As further demonstration of its mechanical toughness, we clamped the wires and the distal end of the device to our motorized strain cycle apparatus and applied a longitudinal strain of 50% for 100 cycles. Even though this test was aggressively applying stress directly to the wires and therefore the bond pads, the increase in total resistance was less than 5% in the piezoresistor circuit. So, whether this was a shift in the piezoresistor or the contact resistance, the wired interface is robust. Lastly, an important feature of the conductive layer was the capability of embedding them in the neutral plane of a relatively thick MEMS device (300 µm). This theoretically allows the device to be bent without adding any stress in the conductor layer. We believe this fabrication solution contributed to the low open-circuit rate of this device during handling and the surgery implantation procedures.

## 3. Conclusion

In this study, we developed an implantable direct bladder interface for strain sensing and bladder muscle stimulation using low-durometer silicone, carbon nanotubes, a Pt-PDMS nanocomposite and a medical-grade super alloy wire. Specifically, single and double-layer CNTs were insulated by silicone (Ecoflex) to form highly-stretchable and sensitive piezoresistive or piezocapacitive sensing devices. We test both 100% uniaxial in benchtop tests and over 90% biaxial strain during *in vivo* tests. The Pt-PDMS electrodes were screen-printed over CNT pads to form a stretchable electrode with the ability to deliver up to 3 mA of current. Additionally, a robust wire connection was integrated onto the device to make an array that has the potential to perform chronically. To our knowledge, this work was the first attempt at evaluating recent developments in stretchable sensors for functional control of a body organ. This device will enable a control system to drive bladder function through embedded logic and closed-loop control. While chronic in vivo validation is required, we hope this work inspires future medical device design for a bladder, cardiac, or gastrointestinal organ interface. For the SCI patient population in particular, having a robust sensor to notify them when their bladder or bowel is approaching full and driving micturition and continence would greatly improve their quality of life.

## 4. Experimental Section

### Device fabrication

To create the device, we utilized both conventional microelectronic fabrication technologies and a series of non-conventional MEMS fabrication and packaging approaches as shown in Figure 2. For simplification, we first detail the fabrication process of the resistive device. Starting from a 4” silicon wafer substrate, we spun on an acetone dissolvable sacrificial layer (Photoresist AZ-5240), followed by a 150-µm thick Ecoflex 00-50 layer (Smooth-on Inc.). Next, single-walled carbon nanotubes (Cheap Tubes Inc. Single Walled-Double Walled Carbon Nanotubes 99, 1-4 nm diameter and 3-30um long) were mixed into IPA at 0.3% wt. and ultrasonicated for 6 hours. Next, the IPA solution with CNT was spray deposited^[59]^ onto the top surface of the spun-on Ecoflex through a shadow mask. Next, Ti/Pt/Au was evaporated subsequently through the second shadow mask on top of the CNT pattern to define the wire bonding pads and the stimulation electrodes. Next, two layers of 100 µm-thick dry-film photoresist (DuPont Riston GM100) were laminated on top and patterned by photolithography. Then, another layer of 150 µm-thick Ecoflex 00-50 was spun-on to complete the encapsulation of the conductors. Therefore, the CNT layer was encapsulated on the neutral plane, which prevented artifact strain induced by device bending or folding during handling or after implantation. Ten microns of the top surface of the whole structure was then etched by reactive ion plasma using SF6/O_2_ with LAM 9400^[60]^. After the top of the dry-film photoresist was exposed, the encapsulation Ecoflex layer was patterned by fully dissolving the dry-film photoresist. For piezocapacitive devices, an additional 5-µm thick insulating layer of Ecoflex 00-30 was spun coated between the lower and upper CNT layers. To access the lower pads of this device, we used a thinner-type dry film photoresist (MM540) and the same RIE-assisted lift-off process to pattern this insulating layer. Medical grade super alloy wires (MP35N PFA Green coated, Fort Wayne Wire) were bonded via silver paste reflow and sealed with additional Ecoflex. In the end, the device perimeter (23 mm by 12 mm) was defined manually with a razor blade but could have been laser cut. Then the device was released by dissolving the sacrificial layer (Fig2. d & e) in acetone. Each sensor had an area of 9.6 mm by 4.8 mm. The stimulation electrode had a 2-mm diameter. Each suture point (patterned with CNT/Au circles) was 1-mm diameter.

Finally, Pt-microparticles and PDMS composite gel was screen-printed into the electrode wells^[51]^. The Platinum microparticle and PDMS composite was prepared by mixing 60% PDMS (Sylgard 184, Dow Corning) with Platinum microparticles (Platinum powder, amorphous, <3-micron, Alfa Aesar) by weight. PDMS was diluted in heptane in a 1:2 ratio to reduce the viscosity. 60mg of diluted PDMS was added to 80mg platinum microparticles in a small centrifugal tube and mixed thoroughly with ultra-sonication. The paste was then printed onto the formed CNT/Au electrodes with a shadow mask and left on a hot plate at 60 °Covernight to ensure complete polymerization. The composite was etched by oxygen plasma to remove the top PDMS so that more platinum can be exposed to deliver charge. The SEM image of electrode surface after preparation is shown in figure S4.

Teflon Pledgets (Ethicon PCP50) were cut to a dimension of 1.5 × 1.5 mm and glued with a small drop of Ecoflex 00-50 to reinforce each suture site for in vivo tests.

### Mechanical Strain Testing

The sensor response to uniaxial strain under physiological conditions (37°C phosphate-buffered saline) was bench tested (Fig. 4,5). Deionized water was added to the solution every 6 hours in order to negate the effect of evaporation. In all tests, the device was clamped horizontally with one end on a linear motor (Thorlabs, KMTS50E) and the other end on a 3D manipulator (Supp. Fig. 3). The devices were then stretched slowly from 0% to 100% strain and repeated for the number of reported cycles. The motor speed was 0.8 mm/second, or about 42 seconds/cycle. Resistance was actively recorded through a voltage divider circuit and sampled (Keithley 2410 and LabVIEW control program) at 54 samples per second.

### In vivo test setup and procedure

Four adult male felines (4.2-5.0 kg)were anesthetized and prepared for non-survival surgery as previously described^[61][62]^. With the animal in a supine position, the bladder was accessed via a laparotomy and slightly elevated above the abdomen to allow for clear visualization. Saline was applied to the bladder regularly to maintain tissue viability. A 3.5-4 Fr dual-lumen catheter was inserted into the bladder via the urethra for fluid infusion and pressure monitoring (Utah Medical DPT-100 pressure transducer, World Precision Instruments TBM4M transducer amplifier, AD Instruments PowerLab data acquisition). A piezoresistive device was sutured on the exposed bladder wall, with the wire end closer to the urethra and the other end positioned near the bladder dome (Fig. 2f). Each pledget on a device was sutured with a single 5-0 nylon knot (661G, Ethicon). Each suture was applied within the bladder wall and did not pass into the bladder lumen. While suturing through the bladder wall may be acceptable for long-term implant validation, we found that during short-term experiments fluid would leak around through-wall sutures at high volumes. The device connections were placed in a simple voltage divider circuit, with a 5 V DC power supply providing current through the device and an in-series resister (~100-350 kΩ). The voltage across the device was measured (AD Instruments PowerLab) and converted to resistance offline. Starting from an empty volume, saline was infused into the bladder at a constant rate (2-12 mL/min) with a syringe pump until leaking from the urethra was observed. After a short hold period (~30 seconds), the saline was withdrawn, and a fill cycle was repeated.

### In vivo strain calculation

The strain of the devices was analyzed from top-view videos recorded during *in vivo* studies using the MATLAB computer vision package. The MATLAB program detected the boundary, corners and centers of the device implanted on a bladder. Biaxial strain was calculated from the change of the real-time device area under strain, which was estimated by the program through a total pixel count inside the device boundary. To analyze the strain in each direction, the program first defined the longitudinal and transverse directions by calculating the normalized vectors from corner to corner. Then, the expansion in either direction was calculated by the dot product of the real-time center-to-center electrode distance and the normalized vector. Finally, the longitudinal and transverse strain were calculated as the percentage device expansion in each direction. Maximum uniaxial and biaxial strain of each device was measured by this approach (Table S1).

### Stimulation-induced bladder contraction experiment

In one experiment, electrical stimulation through the device was performed. Bipolar stimulation (isolated pulse stimulator model 2100, A-M Systems) was applied between two of the electrode contacts on the device. First, the device was laid on the exposed abdominal muscle, after a midline incision in the skin. Low-frequency (3 Hz) and higher frequency (33 Hz) stimulation was applied at 1mA through the device to demonstrate recruitment of somatic muscle. Next, after a midline incision through the abdominal muscle was performed to expose the bladder, the device was placed on the bladder dome for evaluation of different stimulation parameters (33-50Hz, 1.5 to 2.5mA, 3ms pulse width). A urethra catheter placed inside the bladder was connected to a transducer (Utah Medical DPT-100) for monitoring bladder pressure, with an amplifier (World Precision Instruments TBM4M) and data acquisition system (AD Instruments PowerLab) used to store data on a computer.

## Acknowledgements

We thank the NIH for their support through R21EB020811, OT2OD023873, and F31HD094480. We thank other members of the Peripheral Neural Engineering and Urodynamics Lab for their help with animal experiments and device testing including Alex Mundorf, Aileen Ouyang, and Eric Kennedy. We also acknowledge the use of the organic material processing lab area of Prof. Stephen Forrest’s lab, electrochemical characterization facility of Prof. James Weiland’s lab, Lurie Nanofabrication Facility (LNF) at the University of Michigan, assistance from Pilar Herrera-Fierro during the fabrication process and Beomseo Koo during the stimulation electrode characterization experiments.

