## Supplemental and Table 1 to 5 for "Ultra-compliant carbon nanotube stretchable direct bladder interface"

**Support Information**

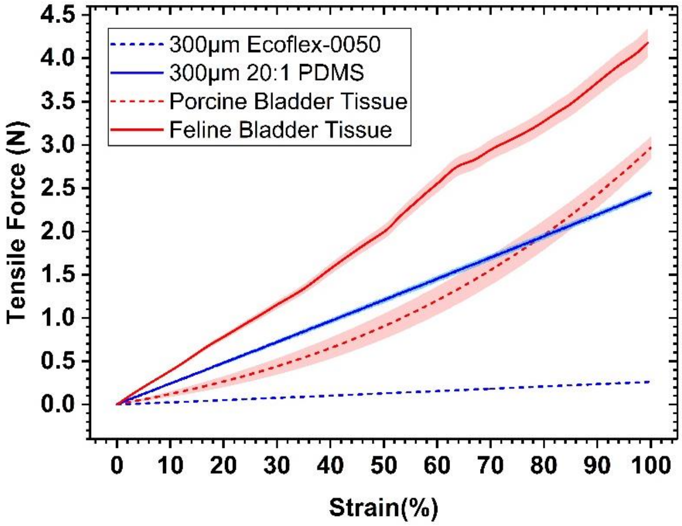


**Figure S1. | Substrate stiffness relative to the bladder wall.** Characterization of the elasticity of three sections of porcine bladder, two sections of feline bladder, Ecoflex, and PDMS. All sections were 1.2cm x 2cm.

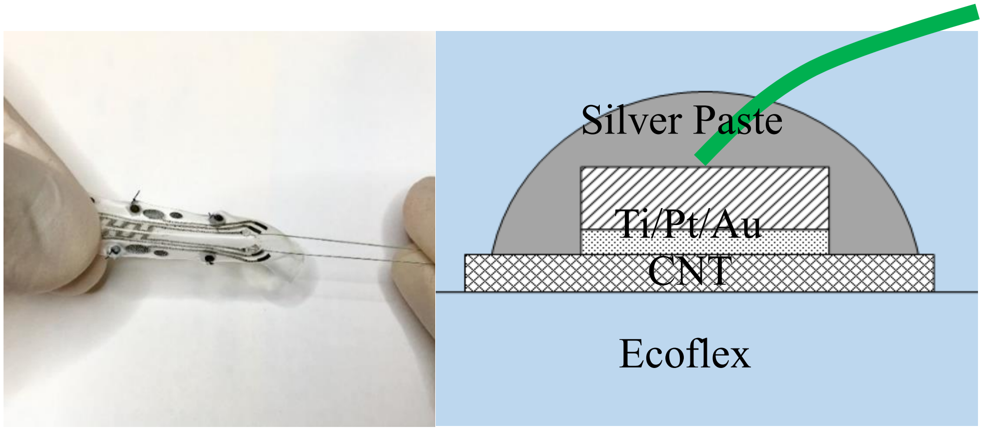


**Figure S2. | Bond pad interface and mechanical robustness.** Left: The wire was pulled directly until the whole device was extended by 50% while no electrical or mechanical failure was observed. Right: The wire bonding interface was made out of a 50nm/100nm/300nm thick Ti/Pt/Au layer and Ag paste. The wire was a medical grade super-alloy MP35N.

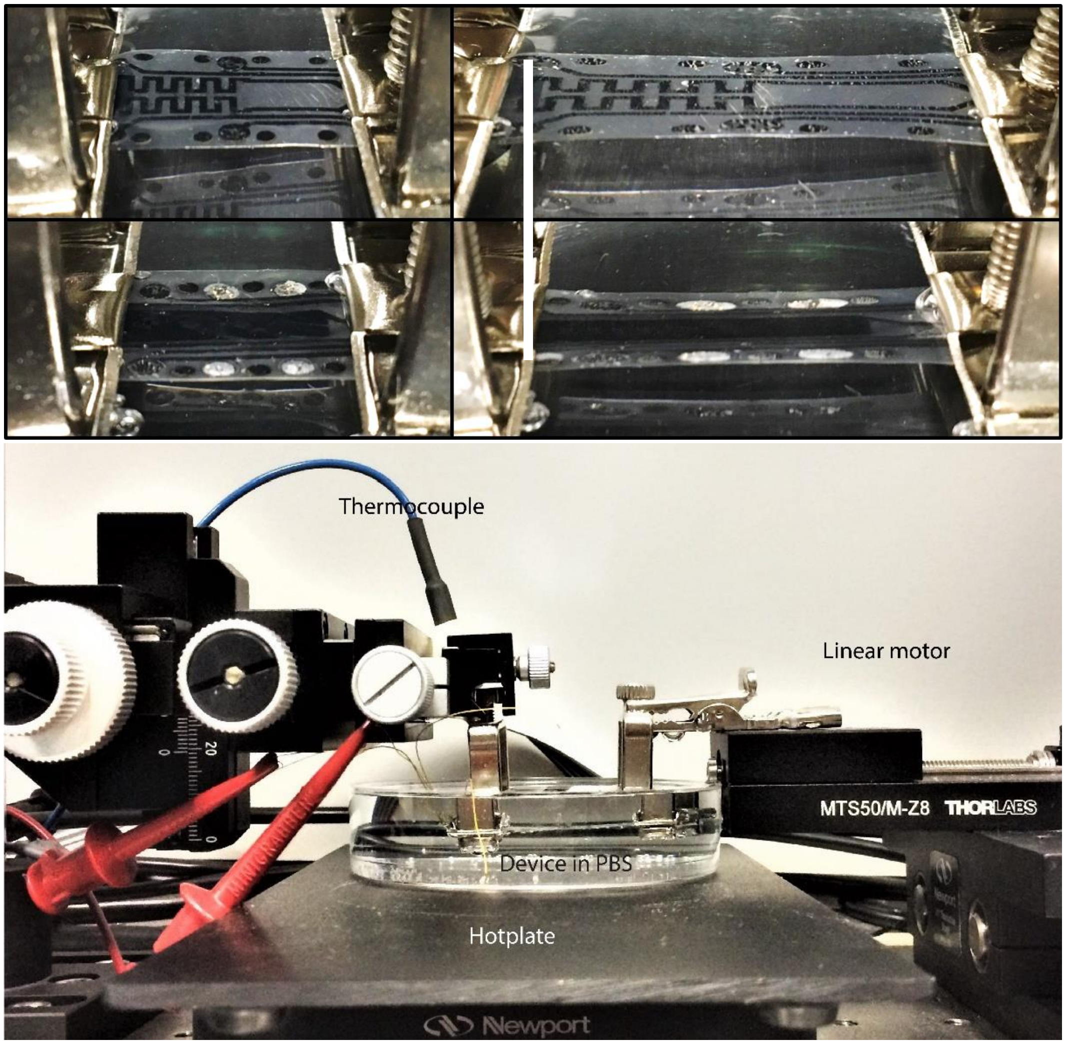


**Figure S3. |Uniaxial test setup. a,** Snapshots of a single device at free state and being stretched to 100% strain. **b,** Photo of the bench-top testing of a device submerged in a phosphate-buffered saline (PBS) bath. The hot plate maintained the bath temperature at 37°C.

| Experiment | Max Strain, Longitudinal ($\Delta L/L$) | Max Strain, Transverse ($\Delta W/W$) | Max Strain, Biaxial Measured ($\Delta S/S$) |
|---|---|---|---|
| **Animal 1** | 44.0% | 31.9% | 82.9% |
| **Animal 2** | 27.8% | 22.2% | 48.8% |
| **Animal 3** | 49.6% | 30.2% | 90.5% |
| **Animal 4** | 41.6% | 28.0% | 71.1% |

**Table S1. | Maximum strain on the device during *in vivo* test.** The biaxial strain was estimated in two ways: calculated from the strain of two orthogonal direction; or measured from device area change by pixel counting.

During bench top testing the transverse dimension was allowed to remain relaxed, therefore based on area change alone, the equivalent biaxial strain range in table above (49 to 90%) was approximately equivalent to a uniaxial strain of 56% to 109%. This was calculated from images of device during bench testing. In an over-filling case, we observed electrical failure at a biaxial strain above 133%, which is approximately equivalent to a uniaxial strain of 160% but that degree of strain was not attempted on the bench.

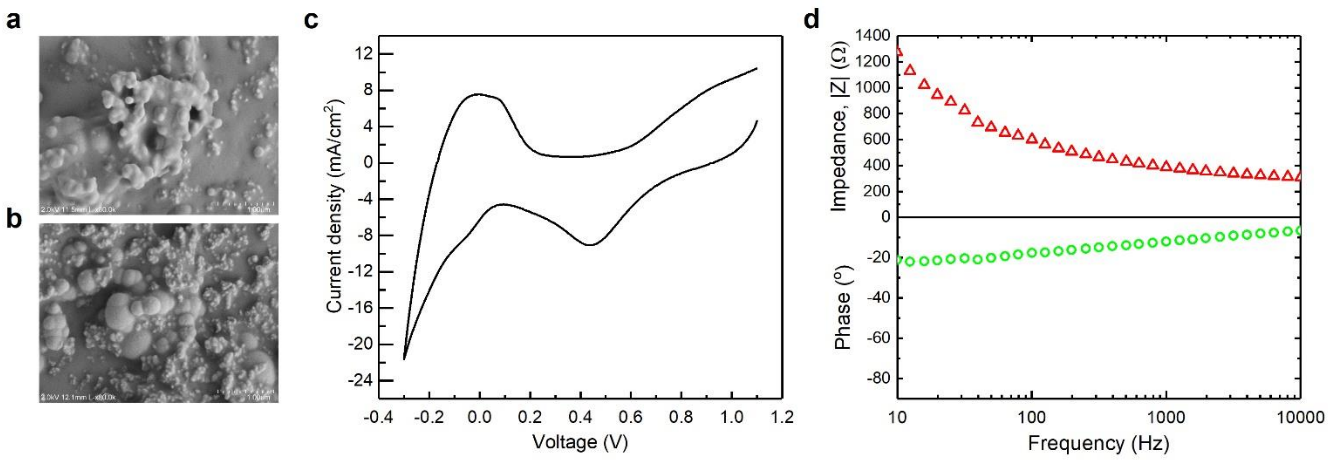


**Figure S4. | Pt/PDMS micro-composite flexible electrode properties. a,** Surface of the Pt/PDMS micro-composite electrode as-printed and **b,** after 20µm plasma etching of PDMS. **c,** Cyclic voltammetry and **d,** impedance spectroscopy characterization of one electrode after the plasma etching surface etching.

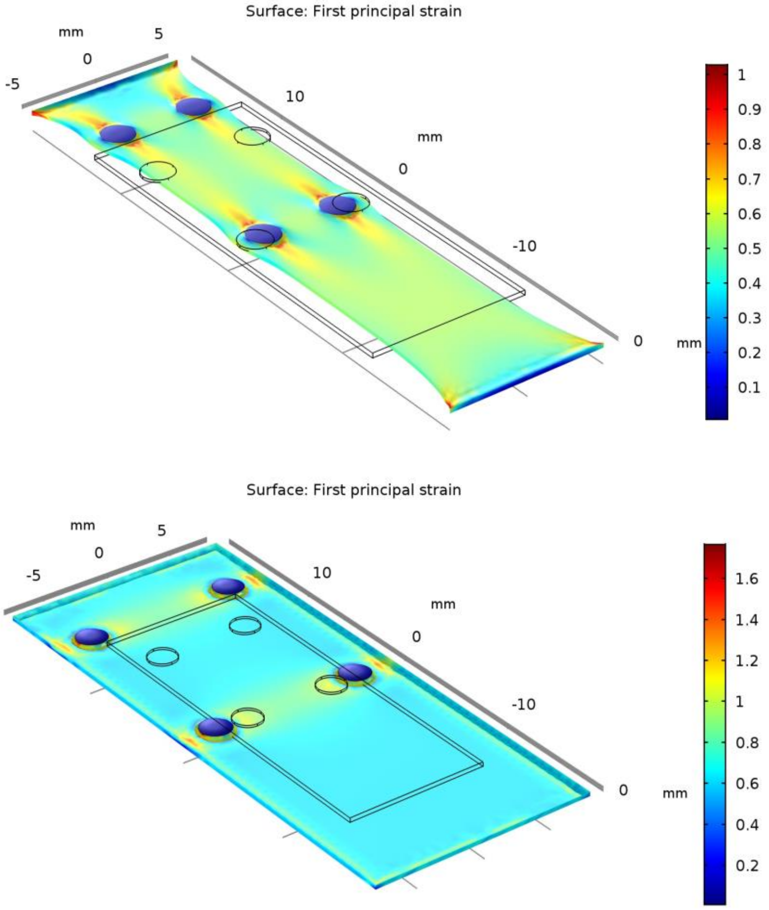


**Figure S5. | Simulated strain distribution around the electrodes.** (top) A 4-electrode device with Pt/PDMS composite had an average strain of only 14% when under uniaxial, 50% longitudinal strain and (bottom) biaxial, 50% longitudinal and 30% transverse strain.

**Supplemental Video 1. | Demonstration of stimulation effects on the abdomen of an anesthetized feline.**
